# Using synthetic semiochemicals to train canines to detect bark beetle-infested trees

**DOI:** 10.1101/262386

**Authors:** Annette Johansson, Göran Birgersson, Fredrik Schlyter

## Abstract

In this proof of concept study, we report the off season training of two detection dogs on a series of synthetic semiochemicals associated with *Ips typographus* pest bark beetle infestations of spruce trees. Scent detection training allowed dogs to discriminate between physiologically-relevant infestation (target) odours, quantified by GC-MS using extracted ion chromatogram to be bio-active at levels of < 10^−4^ ng /15 min or lower, and natural non-target odours that might be encountered in the forest. Detection dogs trained to recognize four different synthetic pheromone compounds in the winter time, well before beetle flight, were able to detect natural infested spruce trees unknown to humans the following summer. The trained detection dogs were able to detect an infested spruce tree from the first hour of bark beetle attack until several weeks after the attack. Trained detection dogs appear to be more efficient than humans in detecting early bark beetle infestations because the canines ability to cover a greater area and by olfaction detect infestations from a far greater distance than can humans. Infested spruce trees could be detected by trained detection dogs out to more than 100 m.

**Key Message:**

- Detection dogs were rapidly trained to locate release of synthetic bark beetle pheromone components
- Synthetics allowed dog training off-season both in laboratory and field
- Dogs trained on synthetics detected naturally target pest insect attacked trees at a distance of more than 100 m.
- The method allows rapid removal of single, first attacked trees before offspring emergence, thus curbing local pest increase and lowering spread of attacks in the landscap

## Introduction

Detection dogs are used to locate many objects including humans, explosives, and illicit drugs (see BROWNE et al. 2006 and references therein; LORENZO et al. 2003). Trained canines have also been used to detect invasive organisms (GOODWIN et al. 2010; HOYER-TOMICZEK et al. 2016) as well as endangered species (reviewed by BEEBE et al. 2016). Canines have also been trained to detect small or cryptic insects such as termites (BROOKS et al. 2003), palm weevils (NAKASH et al. 2000), and bed bugs (PFIESTER et al. 2008; VAIDYANATHAN AND FELDLAUFER 2013) and endangered Coleoptera (MOSCONI et al. 2017). The key benefits of using trained detection dogs are their keen sense of smell (HEPPER AND WELLS 2015), and their ability to cover large areas in a shorter time, when compared to humans (MOSCONI et al. 2017). In most cases, biological material is used for the training (JOHNEN et al. 2013).

The European spruce bark beetle - *Ips typographus* (L.) is one of the most destructive forest pests in Europe (GRÉGOIRE AND EVANS 2004). For forest protection, the rapid detection of bark beetle infestations is required to successfully implement a management strategy that relies upon removing recently infested trees within 2 -3 weeks of attack (SVENSSON 2007). However, human detection generally requires close inspection (≤ 1m) of trees, and is therefore time-consuming, costly, and not always practical. Therefore, detection generally occurs 2-3 months after an infestation in N Europe, when tree crown colour fades and bark falls off. By this time, most bark beetles have left the infested tree and may attack other, non-infested stands. Since a rapidly changing, but specific series of beetle pheromone components and other semiochemicals are present for several weeks after an initial attack, the use of detection dogs may prove a better alternative than human inspection. Upon attacking a tree, male bark beetles secret an aggregation pheromone, consisting of a blend of 2-methyl-3-buten-2-ol and *cis*-verbenol (BIRGERSSON et al. 1984). A few days later, an inhibitory signal (consisting mainly of ipsdienol) is emitted when bark beetle females have begun laying eggs (BIRGERSSON et al. 1984; SCHLYTER et al. 1987). After the first week, an additional chemical cue, indicating that the infested tree is fully utilized and competition is high, is evident. This semiochemical, verbenone, is an oxygenation product by the beetle and by the interaction of fungi and bacteria with damaged tree phloem (LEUFVÉN AND BIRGERSSON 1987; SCHLYTER et al. 1989).

In this proof of concept study, we report the laboratory training of two detection dogs on a series of synthetic semiochemicals associated with bark beetle infestations, and the ability of these trained dogs to later detect and locate bark beetle infested trees in the field. Since the semiochemical profile of attacked trees changes rapidly in both the quality and quantity of semiochemicals released, we chose to use several synthetic chemical compounds as representative stimuli in our canine training. We were also interested in determining if dogs trained on synthetic pheromones in the winter months could later locate infested trees in the summer months. Finally, we wanted to determine if a trained dog can detect natural infestations from distances (10 to 100 times) further away than a human.

## Materials and methods

### Canines

Two dogs, owned by SnifferDogs Sweden (Hjortsberga, Sweden), were used in this study. Dog A was a nine-year-old female German shepherd that was previously trained as a search and rescue dog for humans. Dog B was a one-year-old female Belgian shepherd (Malinois) that had only basic obedience training, and had no previous formal detection expertise.

### Chemicals

Synthetic bark beetle pheromones used in this study included methylbutenol (2-methyl-3-buten-2-ol; Acros Organics, Gothenburg, Sweden), 4S-*cis*-verbenol (Borregard, Sarpsborg, Norway), and (*S*)-ipsdienol (Bedoukian, Danbury Connecticut, USA). Synthetic verbenone, a bark beetle pheromone and a product of the host tree was obtained from Fluka (Sigma-Aldrich, Stockholm, Sweden). Other chemicals used in the study were obtained from our chemical stocks (see ANDERSSON et al. 2012).

Each pheromone component was stored separately in separate jars of glass, to avoid cross contamination of odours. In each jar of glass a cotton pad (ICA Basic Bomullsrondeller, Netherlands) was placed in the bottom and a small amount of each semiochemical were dropped on to the cotton pad (10 μl methylbutenol, ≈10 mg *cis*-verbenol, 1μl ipsdienol, or 10 μl verbenone). The glass jars were then filled with cotton pads, and so molecules in gas phase of each component passed passively by aeration transfer in the closed jar, via adsorption of the odour to the pads placed above (HUDSON-HOLNESS AND FURTON 2010). The glass jars were stored in a freezer (≈ –18 °C).

For determination of release rates by GC-MS and dog training response (Fig 1, Table 1) we always used the cotton pads from the top in each glass jar. The last five cotton pads in the glass jars we never used but filled up with new pads when needed. A cotton pad holding the semiochemical (HUDSON-HOLNESS AND FURTON 2010) was placed in a stainless steel tin (5 cm dia.) with perforated lids (“Ströare”, Biltema®, Helsingborg, Sweden). For ordinary dog training, we replaced the cotton pads each day. For release rates by GC-MS and dog training response study we prepared 5 steel tins of each synthetic pheromone at the same time. These tins were stored in room temperature (circa + 20 °C). Release rates were determined using odour collections similar to Zhang et al. (2000). An inverted glass funnel (5 cm dia.) was placed above the steel tin and air was drawn through a column packed with Porapak® Q (25 mg mesh 60-80; in a Teflon tube 3 mm i.d.) at 100ml/min at 15 min intervals. Compounds were eluted from the column with 400 μl pentane (Sigma-Aldrich, Steinheim, Germany) into a 400 μl insert placed in a 2 ml screw top vial (Agilent Technologies, Böblingen, Germany), and 1 mg heptyl acetate was added as internal standard. Aeration extracts were analysed by gas chromatography-mass spectrometry (GC-MS; Agilent 6890-5975, Agilent Technologies, Santa Clara, CA, USA) with techniques previously reported (BIRGERSSON et al. 1984). Quantifications were based on extracted ion chromatograms of prominent fragments for each tested compound and the internal quantification standard, respectively. The limit of quantification (LOQ) in the analytical procedure was < 0.1 ng/min.

**Fig 1.**
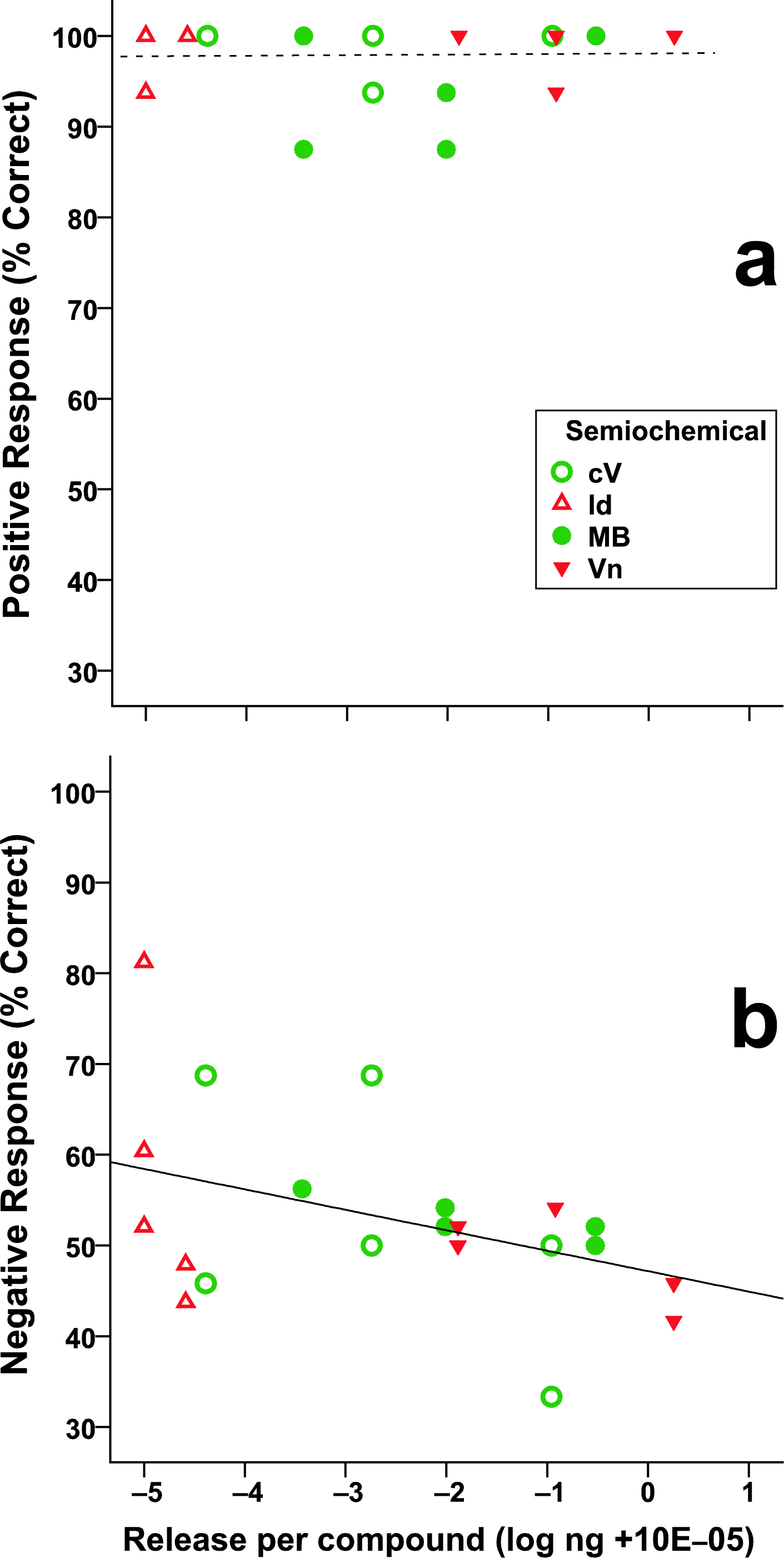
Correct responses in relation to estimated stimuli evaporation rates per 15 min. **a)** Correct positive responses to stimuli (compounds) with known total release over 3 days of testing. No effect of time for all stimuli joined (*r*^2^≈ 0). Correct negative responses to stimuli with known total release over 3 days of testing. Weak effect of time for all stimuli joined (*r*^2^= 0.18). Separately, MB shows the strongest effect (*r*^2^= 0.85) together with Vn (*r*^2^= 0.48). Chemical data from Table 1. The responses of the two dogs are pooled here, separate data in Table 2. Semiochemical acronyms: MB) Methylbutenol, cV) 4*S*-*cis*-Verbenol, Id) Ipsdienol, Vn) (–)-Verbenone.

### Laboratory tests

Initially, dog A was introduced to the bark beetle pheromones using the synthetic odour from a commercial dispenser, ETOpheron ® (Pheronova AG, Switzerland), which is used in bark beetle monitoring traps. Because of the dispensers construction of fabric with a plastic shell it was not 100% sure that the dog learned the scent of the pheromone components as the target odour or if it learned any other odour of the dispenser materials. It is easy to inadvertently train a dog to detect an unexpected or impure source when attempting to train to a pure compound. To be sure that the dog learned the right odours we subsequently trained the dog on pure synthetic semiochemicals applied to cotton pads. Non-target odours, that could disturb search, were also used in the training and consisted of items found in a forest setting such as vegetation odours from spruce needles, cones, resin, bark, moss, and animal odours (i.e. scent from feathers, fur, hoofs and faeces). All non-target (disturbance) odours were collected in the forest, or donated by local hunters (fur and hoofs from moose, deer, and boar). Both target and non-target odours were stored in jars of glass and transferred by aeration to cotton pads to ensure that the background odour of cotton was present in both target and disturbance odours.

The training platform used here (2D illustrations in Figure ESM_1 and video in ESM_4_V1), was developed by Stig Meier Berg and Geir Kojedal, Spesialsøk, Selbu, Norway, based on an idea from Hundcampus, Hällefors, Sweden (FISCHER-TENHAGEN et al. 2011). It is designed to let the dog work independently, to minimize the cues from the handler and to be easily manoeuvred by the handler creating a more effective learning situation with a high rate of opportunities to reward the dog for desired behaviour. Disturbance odours were presented together with one or several semiochemical stimuli in a movable tray with 7 positions (Figure in ESM_2).

For evaluation of the dog detection performance with decreasing amounts of odour molecules over time nine trials were conducted to evaluate the dogs’ identification performance with each synthetic semiochemical. For this trials we used 4 of the prepared 5 steel tins containing cotton pads with synthetic semiochemical. Since the trials were conducted over several days (1 hour through 84 hours after the cotton pads being placed in the tins and stored in room temperature) we used a new tin every day. This was done to make sure that the tins weren’t contaminated with any other scents such as odour from the dogs. Every trial session lasted for approximately one minute (50-70 seconds).

To compare different stimuli linear layouts, mixing target and non-target scent, on the movable tray, the two dogs were tested in three trials with each stimuli layout (ESM_2).

### Outdoor tests

To train the dogs to pinpoint the target odour source outdoors, pieces of the cotton pads containing synthetic pheromone odours as those used for platform training were hidden in cracks of the bark of several species of trees. The cotton pieces were placed in the height of the nose of the dogs and the dogs were shown where to sniff for the target (video ESM_4_V2). When the dog found the cotton piece holding the target odour, it was immediately rewarded by the sound of the clicker and a piece of food delivered between its nose and the odour source. Several pieces of cotton with either target or non-target odours, were put in cracks of the bark in a small area (30×30 cm) to ensure the dogs did not use visual cues for close-range target location. When the dogs were able to consistently (∼ 100%) locate the pads, the dogs were gradually sent from longer distances to locate the tree with the cotton pad, allowing the dog to detect decreasing amounts of target molecules and to follow the odour to its source.

Dogs were trained in the winter under a variety of weather conditions (*e.g.* rain, snow, sun). Training trials using synthetic odour were conducted on average once a week during 2009 and 2010. The temperature ranged from 2 to 28 °C. The handler determined the search strategy to best cover the assigned area based on wind conditions and terrain. These protocols were employed to simulate future practical field survey conditions.

A proof-of-concept test, evaluating the detection by dogs of spruces that were known to be recently attacked by bark beetles, was conducted at the Nature Reserve of Notteryd (near Växjö, Småland, Sweden). The area consisted of wind-felled trees and standing healthy spruces. In the spring of 2009, 95% of all spruces in the reserve were already killed by bark beetles. The remaining spruces that were still alive stood together in clusters of 10-15 trees. We felt these circumstances made this particular Nature Reserve an optimal area to first try the dogs on natural attacks. Another series of tests were conducted at a production-forest in Nottebäck, also near Växjö, Småland, Sweden, with the permission of the owner of the forest. The dog team consisted of one dog (dog A) working off-leash and one handler, and searched three different areas in the production-forest attacked by bark beetles in previous years. The handler had knowledge of the location of former attacks, but no information if there were any new attacks. The dog and handler searched each area with no time-limit. The handler determined their search strategy to best cover the assigned area based on wind and terrain. These protocols were employed to simulate expected future practical field survey conditions.

Dog and handler movements were recorded using global positioning systems (GPS) in 5-second intervals in all field trials. These data allowed identification of the point at which the dogs lifted their nose up in the air and made a sudden change in direction of travel and moved directly towards an infested spruce (video of search ESM_4_V3). The GPS units used in the study were Garmin Astro® 220 Nordic handset and Garmin DC30 dog collar (Garmin Corporation, Taiwan). The map used in the handset was Garmin “Friluftskartan Pro V2 Götaland”. The data from the GPS-unit were transferred to a PC with Garmin’s software MapSource. Using the measuring tool we could measure the distance from where a track from the dog changed direction to the waypoint where the dog alerted on an infested spruce.

#### Field trial - detection distance from natural sources

We used 20 different areas, whereof 10 were located in nature reserves and 10 in production forests, with permission of the owners and from The Swedish Forest Agency in Växjö in 2010. All areas were 2 - 4 ha and tests were done in three different set-ups; a) 10 search areas with location of infestations known by handler b) 5 areas with location of infestations known by the forest manger, and c) 5 areas with location of infestations unknown, but were considered as risk areas with bark beetles infestations previous seasons.

To design the best search strategy for long distance detection, based on wind, terrain and the location of the attacks, the dog handler had prior knowledge of attacks in the 10 first areas. To estimate if the dog handler might involuntarily que the dog to an odour source (an infested tree), the dog handler was not allowed prior knowledge of attacks in the 10 latter areas.

## Results

### Laboratory tests

The two dogs were successfully trained to recognize the four different synthetic semiochemical compounds on the educational scent platform. Both dogs learned to recognize a new target scent in just one training trial, similar to Johnston (1999). In that time, the dogs managed to sample the tins for target odour about 30 times on average (video ESM_4_V1). Occasionally, the dogs reacted with an increased interest when a new non-target disturbance odour was presented. When this happened, the handler stood silent and just waited until the dog stopped investigating the new non-target odour and, if the dog did not continue to search by itself, gave the dog a new command to start sampling the other tins again. After a few encounters with the new non-target odour the dogs’ interest decreased since they learned that there would not be any reward for that particular odour. Even though the dogs appeared to alert on new, disturbance odours (mostly edible items like cookies and chips or scents from other animals) the handler did not record such behaviour as an alert. When alerting on a target odour, both dogs stopped sampling, and waited for their reward, in contrast to increased sampling a tin in order to investigate a disturbance odour.

### Chemical stimuli strength

Chemical quantification by odour collection and GC-MS was routine with a limit of quantification (LOQ) of < 0.1 ng/min. However, two days later, we found that most stimuli titres, still well biologically active, decreased to below the LOQ. Using estimates based upon linear regression, chemical data indicates that by the third day some compounds were very close to zero (Table 1).

The dogs responded to estimated doses of 10^−4^ ng/15min releases or less. The four different semiochemicals were learned equally well, and responses to sub-picogram release rates of stimuli aged up to 3.5 days remained stable (Table in ESM_3).

### Biological responses

The responses of both dogs to target odours are summarized per target scent in Table 2. The dogs achieved a mean of 99% correct indications; 1% of the incorrect indications were either false positive (alerting to a non-target odour; dog A) or false negatives in the beginning of a trial session (dog B) (Table 2). None of the dogs sampled all tins in every repetition. In each repetition four tins were presented, but the trainer could never know where the dog would start searching or in which direction it would continue its search. The only dispenser tin always sampled was the tin holding the target scent. This explains the high success rate of 99% for correct positives for the target scent (at which tin the search will stop), but the much lower rate, 55% for the correct negatives with direct sampling of empty tins before finding the target scent.

To increase the dogs sampling of all presented tins, we tried the dogs in different kinds of stimuli layouts with zero to three different target odours presented in the same trial, Figure A in ESM_2. The only clear effect was for the layout with no target scent, where response decreased with time, Figure B in ESM_2

Interestingly, over the >3 days of testing combined with chemical sampling, the correct responses remained consistently high irrespective of substance (Table 2). The positive responses showed no decline with estimated chemical stimuli levels (Fig 1A), indicating that stimuli levels were above animal detection limit for the period. The correct negative responses (no alert to disturbance odours) declined with the estimated stimuli strength since the dogs learned that these odours weren’t going to be rewarded (Fig 1B).

### Outdoor tests

During off-season training, dogs were introduced to cotton pads initially placed at nose height in the cracks of the bark in different kind of trees (Video ESM_4_V2). Dog A, previously trained as a search and rescue dog, just needed to come into contact with one of the new learned target odours to expect a reward hence follow it to the source and pinpoint it to its handler. Dog A also alerted the found target source by barking. Probably because this was the trained alert when locating a hidden human as a search and rescue dog. Dog B, however, which had no previous search training, had to learn how to follow the odour plume to the source. Dog B was not trained to perform any other alert than pinpointing the source of the target odour. This dog did not know any other way to receive its reward but putting its nose on the target source. The target source became a button to push to get its reward.

Bark beetle activity usually began at the end of April, when the temperature increased to over 20 °C, allowing us to test the dogs’ ability to detect natural pheromone from attacking spruce bark beetles. Dog A successfully found the first spruce that was under attack, on the first day. The spruce in question showed no signs of the attack at first sight, but further inspection at close range revealed that the first bark beetles were drilling their way in to the spruce bark and the sound of their drilling could also be heard. This finding was crucial in demonstrating that it is possible to train a dog on a synthetic odour and subsequently showing that it will alert to the natural odour under field conditions. All training and detection beetle in the Nature Reserve was terminated at the end of May when so many spruces were under bark beetle attack that the smell from the attacked trees became obvious even for the human nose.

Both dogs were also successful in locating sparser attacks in production forest stands, where attacks were neither known to the dog handler nor the forest manager. In the first area searched, dog A detected and alerted to a single, wind-felled spruce that had been infested by bark beetles. In the second area searched, the same dog found seven infested standing spruces. Five of them stood together in a cluster among old attacks. Two were located in a felling edge.

In the third area, the dog detected five infested spruces, both standing and wind-felled. In this area all the spruces were located near a felling edge by a clear-felled area where felled trap-trees were placed. The dog started its search with detecting and alerting on the synthetic pheromones from the trap-trees. When sent to continue its search the dog detected, recognized, followed, and alerted on the natural pheromones emitted from the bark beetles in standing trees (as shown in video ESM_4_V4).

The handler observed by GPS a majority of successfully located sources of natural pheromone to be detected within 50 m, but both dogs located sources in a behavioural sequence over a range of 50 - 100 m (Fig 2A). No differences in detection distance by GPS could be seen among areas with attacks (10) known or unknown (10) to the handler. Later analysis of the GPS-tracks showed several occasions where the more experienced dog A changed direction and was able to detect the pheromones from bark beetle attacked trees at a distance of over 100 m from the source and follow it to the source (Fig 2B). In the 20 areas visited, the dogs found in total 193 trees infested by bark beetles, in 77 different groups of attacked trees.

**Fig 2.**
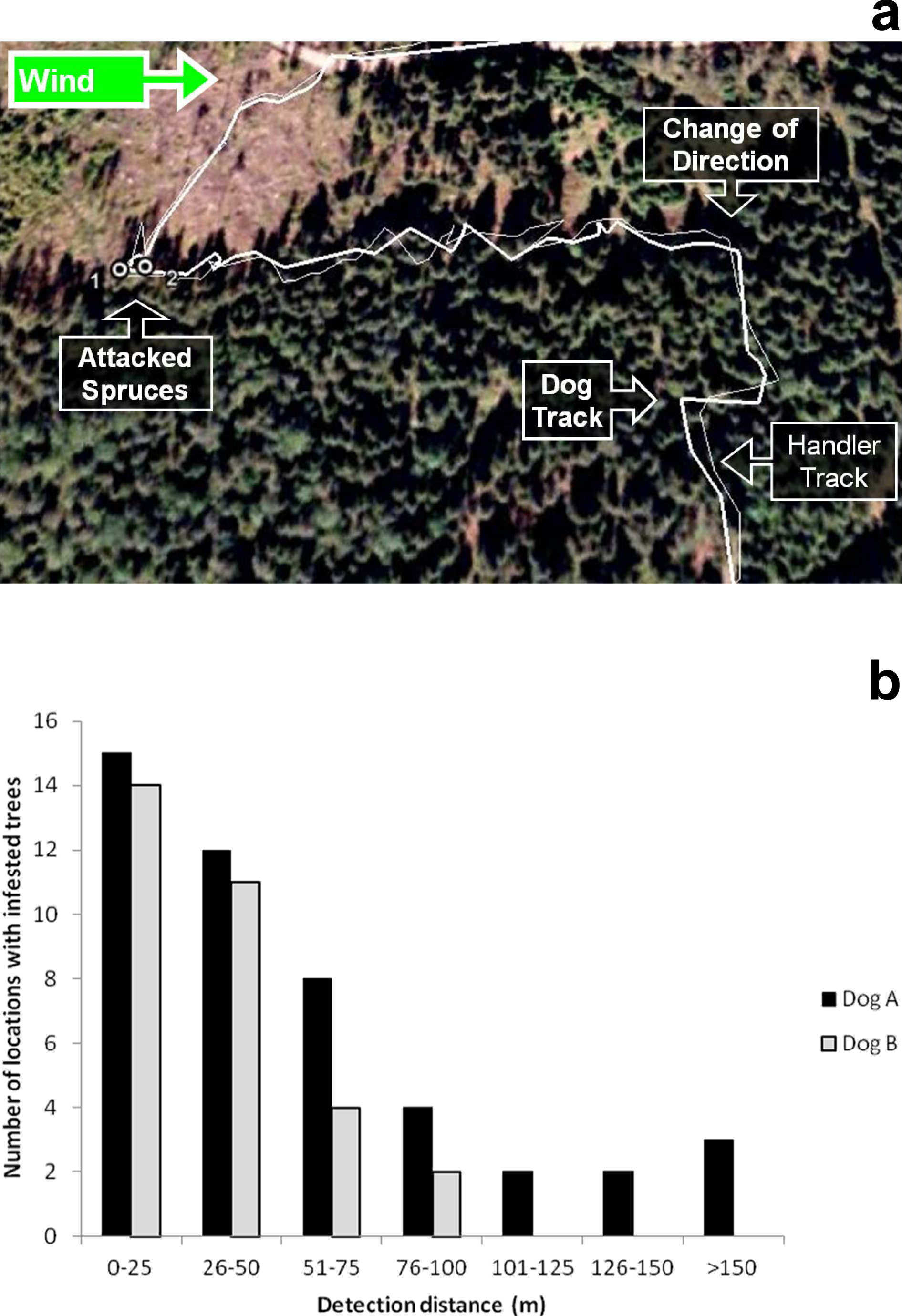
Field GPS tracks and detection distances. **a)**Tracks from an example of handler and search dog finding an unknown mass-attacked single tree. GPS-units tracks shown with Google Earth on a background satellite image over the area. Distance from Change of Direction to the attacked trees = 157 m. Maps, aerial photos/satellite images: Copyright/Lantmäteriet, Sweden, consent #: I2011/0096. (See GPS unit with tracks in video ESM_4_V3.) b)Detection distances recorded from GPS tracks when locating natural bark beetle mass-attacks unknown to handler.

## Discussion

Training canines to detect bark beetle-infested trees poses some important limitations, including the relatively short season available for using trees at various stages of attack, as well the risk of inducing a full-blown tree attack by placing pheromone for training purpose on a host tree during the actual beetle flight period. While it is probably possible to train a detection dog to locate spruces that have been attacked by bark beetles by just letting the dog sniff an attacked spruce and reward the dog, such a “natural” method will not teach a dog to recognize the different kinds of semiochemicals the bark beetle releases over the course of an attack. Therefore, we chose to train the dogs to recognize a series of synthetic pheromone compounds and using an indoor training platform. In this study, we demonstrate that canines trained on synthetic bark beetle pheromone compounds at low (sub-picogram) levels, indoors, can later recognize naturally-produced pheromone over long distances, outdoors.

Additionally, by using synthetic sources of the bark beetle pheromone in the laboratory, it is possible to train dogs off-season long before the bark beetles start their flight period in the field, and the dog handler has control over which odours the dog learns, one at a time and at very low concentrations. The indoor training of canines also has the benefit in that other environmental distractions are minimized, thereby allowing the dogs to concentrate on and learn the target odours.

In the field, detection dogs that work over large areas (“off-leash”) can often be seen lifting their nose up in the air and then make a sudden change in direction of travel. This likely occurs when the dog enters an area with a detectable odour (a plume) that the dog identifies as its trained target odour. While the odour plume structure in a field setting, where the plume shape, size, and persistence is highly dynamic, cannot be easily delineated by chemical means due to the very low titres present in open air (MURLIS et al. 2000; RIFFELL et al. 2008), it can, at least, be observed through olfactory-behavioural responses of dogs to target odour plumes. In our study, a trained detection dog could detect an infested spruce tree from a distance of 150 m, which is farther away than that estimated for bark beetles (*Ips typographus*) responding to beetle pheromone dispensers (SCHLYTER 1992).

In training dogs to detect bed bugs (*Cimex lectularius* L.), the dog usually searches (either “on-leash” or “off-leash”) the entire room – often several times – before alerting on a bed bug. In this case, the dog-handler interaction is paramount owing to the vastly different scales of indoor room searches (1 - 10 m) compared to free-ranging forest searches (10 - 500 m). Issues surrounding a close interaction between dog and handler have been reported (LIT et al. 2011), though during our large scale, forest searches these issues would be minimal, at best. In a study of canines involved in bed bug detection, a high degree of false positives and low true positives were found (COOPER et al. 2014).

Little, if any, studies can be found using pure, known synthetic samples for canine detection purposes (JOHNEN et al. 2013). However, it is clear that canines can show a dose-response to relatively low (but quantitatively unknown) doses (KRESTEL et al. 1984; POLGÁR et al. 2016; WALKER et al. 2006). Our levels of correct positives (sensitivity) and correct negatives (specificity) appear high, compared to the seven studies recently reviewed that provided such data (JOHNEN et al. 2013). Hitherto, no quantitative data exist on chemical strength during dog training in the open literature in spite of some early attempts (KRESTEL et al. 1984; WALKER et al. 2006). No doubt, the dearth of chemical data is due to low thresholds for dog response to volatiles. While our data are novel, we must admit that our empirical data spans only a part of the tested range of stimuli diminution over time, mainly the one-day-old dispenser material, and we had to rely on estimates from linear regression for the older material with lower releases. Still, our estimates appear to be the best so far documented.

The “search-and-pick” method of detection and removal of bark beetle-infested trees within 2-3 weeks of attack (SVENSSON 2007) often fails because of the short time frame involved. Trees were often not cut and removed from the forest until weeks or months later (LÅNGSTRÖM AND BJÖRKLUND 2010), long after beetles had moved to attack other trees. Finding spruces in an early stage of attack is also significant for the timber value, due to a blue stain fungi the beetle introduces into the newly-attacked trees (KIRISITS 2004).

Since the pheromone blends used by the bark beetles for intraspecific communication vary in strength and composition over time, we importantly observed that the dogs could detect all of the substances on which they were trained; therefore, it makes no difference which semiochemical composition the bark beetles in an infested tree is currently emitting. Thus, a trained dog will detect and follow any of the odours, alone or in blends, to the source and alert the dog handler. While it is possible that the dog may learn additional odours that may occur when a spruce is under attack (BIRGERSSON AND BERGSTRÖM 1989; SCHIEBE et al. 2012) any conclusions to this effect would be speculation on our part.

In our study, searches would often be conducted in colder and wetter periods, in-between the short warm-weather swarming periods of the beetle. It would be interesting and of practical relevance to know more precisely how different weather conditions may affect the dogs’ ability to search a larger area.

In view of the large number of pheromones identified to date from moths, beetles and other pests (> 1 000) (EL-SAYED 2017), it would seem feasible to start training of detection dogs for many pest management systems.

In summary, this is the first report of using synthetic pheromone compounds at known titres to train detection dogs to detect and locate living animals in the field. Dogs could detect and locate the source of pest insect infestations at a distance of over 100 m or more. The use of detection dogs for early detection of bark beetle infestations could contribute to better forest protection. We suggest that, in general, use of stimuli that are biochemically well-defined in both quality and quantity appears to hold promise for both better practise and science in detection dog training to biological objects, such as cryptic pests.

## Miscellaneous information

### Funding

See information in Acknowledgements.

### Ethical approval

Both dogs participating in this study were private owned working dogs and handled by their owners. According to Swedish legislation no part of this study included abuse to an animal at the time of study.

### Author contributions

AJ and FS designed research; AJ and GB collected data; AJ, GB, and FS analyzed data; all authors contributed to the writing process. All authors read and approved the submitted version.

#### Acknowledgements

We thank Drs. M. Andersson and A. Bonaventura, and Prof O. Anderbrant, as well as Drs. Veronique Martel and Krista Ryall and Mr T. Gustafsson for valuable comments on earlier versions. Special thanks to go to Dr M. Feldlaufer (USDA, Beltsville) for extensive suggestions on content and language. The authors thank Anton Holmström at The Swedish Forest Agency, Växjö and the private foresters for granting us access to field trial and search areas in nature reserves and production forests. This research was funded by two grants from “The Södra Foundation for Research, Development and Education”, Växjö, Sweden to AJ. FS & GB was supported by the Linnaeus programme ’Insect Chemical Ecology, Ethology and Evolution’ (IC-E^3^, #217-2006-1750) at SLU and later to FS (’Rapid olfactory detection of insect and fungal damage in forests’, #2013-1583) both from “The Swedish Research Council Formas”. FS was further supported by EXTEMIT-K project financed by OP RDE at Czech University of Life Sciences Prague (CZ.02.1.01/0.0/0.0/15-003/0000433). Funding sources had no involvement in study design, data collection/interpretation or writing/submission of this report.

## Supporting Information

**ESM_1** Fig. Educational scent platform. (PDF)

**ESM_2** Fig. Training platform stimuli layout and decline in response to no target scent. (PDF)

**ESM_3** Table. Evaluation of the dog detection performance as number of indications with decreasing amounts of scent molecules over time. (PDF)

**ESM_4_V1** Video. Educational scent platform in operation. (AVI)

**ESM_4_V2** Video. Placement of cotton scent pad and the location of the scent by dog on a pine (a non-host tree of the beetle). (AVI)

**ESM_4_V3** Video. The search, GPS tracking, and location of natural attacks. (AVI)

**ESM_4_V4** Video. The search, location of two adjacent natural attacks, and rewarding. (AVI) (The four videos can be accessed here as well: V1 to V4 Suppl material.)

---

Video stills

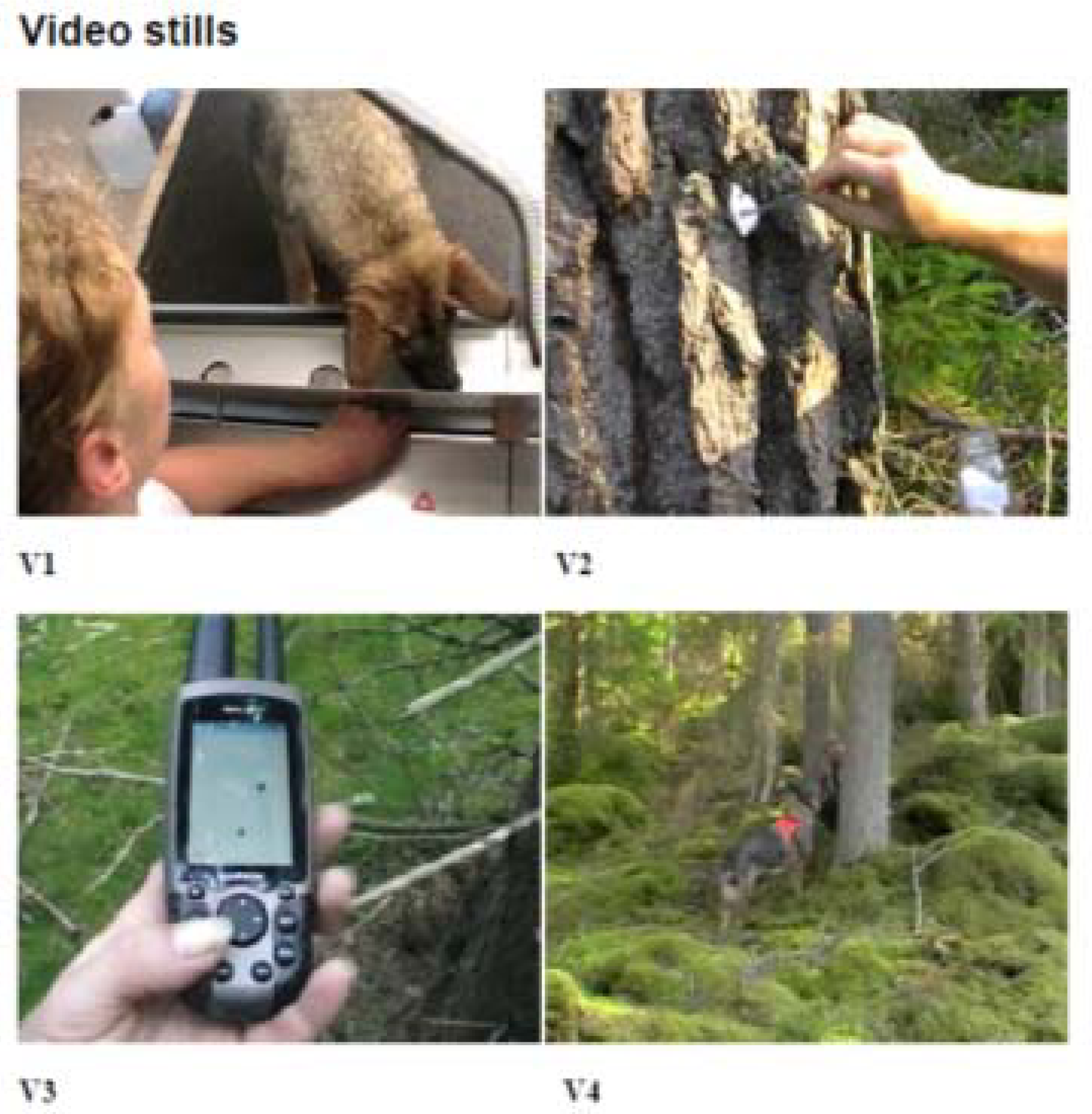

